## Supplementary Figures for "An improved epigenetic counter to track mitotic age in normal and precancerous tissues"

### Supplementary Information for "An improved epigenetic counter to track mitotic age in normal and precancerous tissues"

#### SUPPLEMENTARY FIGURES

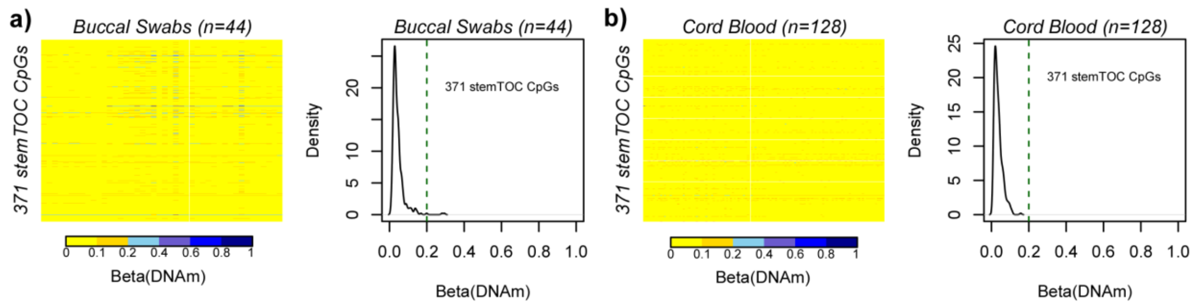


**Supplementary Figure 1: stemTOC CpGs retain ultra-low DNAm levels in neonatal tissues. a)** Heatmap of DNAm values (EPIC) of the 371 stemTOC CpGs in 44 buccal swabs from newborns [1]. Density distribution displays the average DNAm of the 371 CpGs across the buccal swabs. **b)** As a) but for a cord blood Illumina 450k dataset of 128 samples [2].

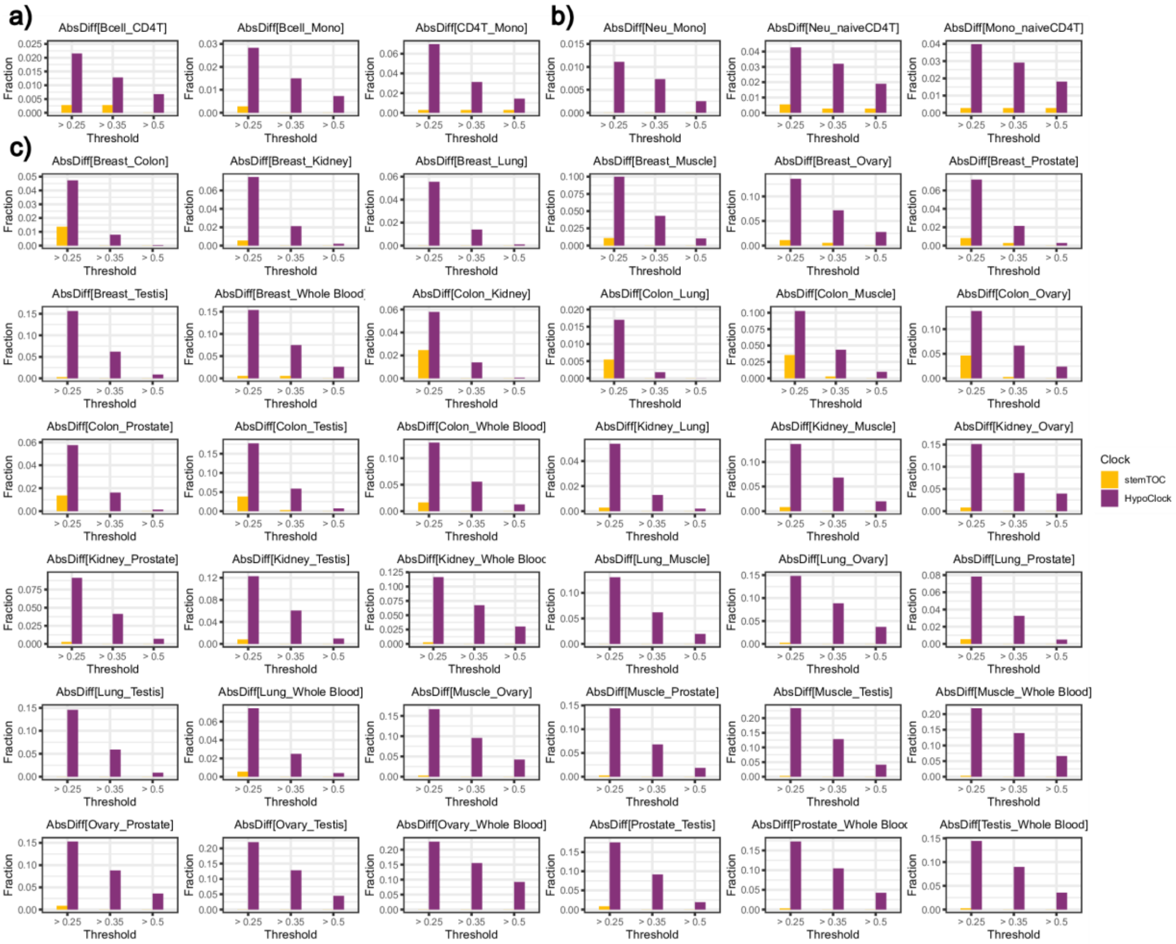


**Supplementary Figure 2: stemTOC CpGs do not display cell-type specific DNAm differences between age-matched sorted immune cells and between eGTEX tissue-types. a)** Barplots depicting the fraction of stemTOC (orange) and HypoClock (purple) CpGs that display absolute differences in DNAm greater than 0.25, 0.35 and 0.5, between age-matched sorted immune-cell types (B-cells n=100, Monocytes n=104 and CD4T-cells n=98 from Paul et al [3]). **b)** As a) but for the matched sorted immune-cell types from BLUEPRINT (Monocytes n=139, Neutrophils n=139, naïve-CD4T-cells n=139) [4]. **c)** As a) but for the age-matched tissues from eGTEX project: breast mammary tissue (n=52), colon transverse (n=224), kidney cortex (n=50), lung (n=223), skeletal muscle (n=47), ovary (n=164), prostate (n=123), testis (n=50) and whole blood (n=54)) [5].

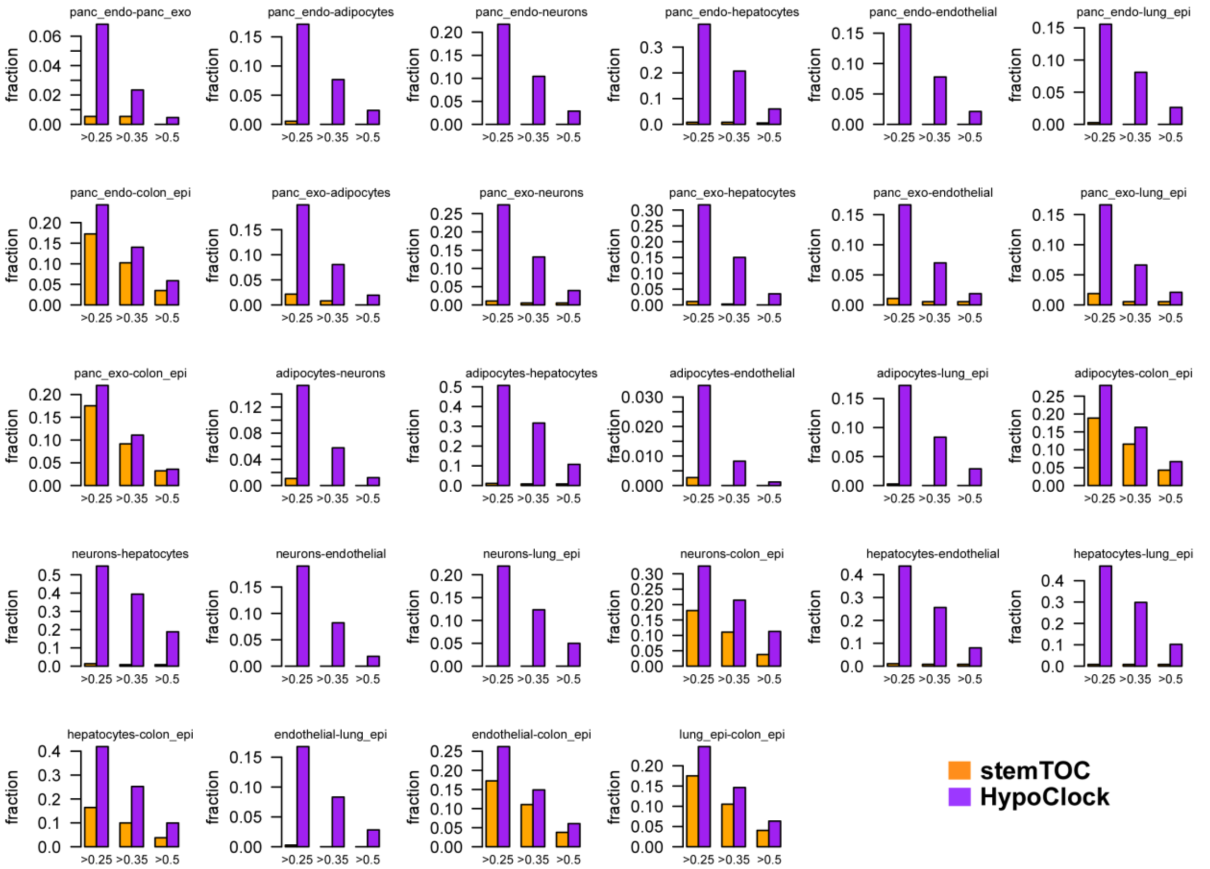


**Supplementary Figure 3: stemTOC CpGs do not display cell-type specific DNAm differences between age-matched sorted cells from Moss et al DNAm-atlas.** Barplots depicting the fraction of stemTOC (orange) and HypoClock (purple) CpGs that display absolute differences in DNAm greater than 0.25, 0.35 and 0.5, between age-matched sorted cell types from Moss et al [6]: sorted pancreatic beta cells (n=4), pancreatic ductal (n=3), pancreatic acinar (n=3), adipocytes (n=3), hepatocytes (n=3), cortical neurons (n=3), leukocyte (n=1), lung-epithelial (n=3), colon-epithelial (n=3) and vascular endothelial (n=2).

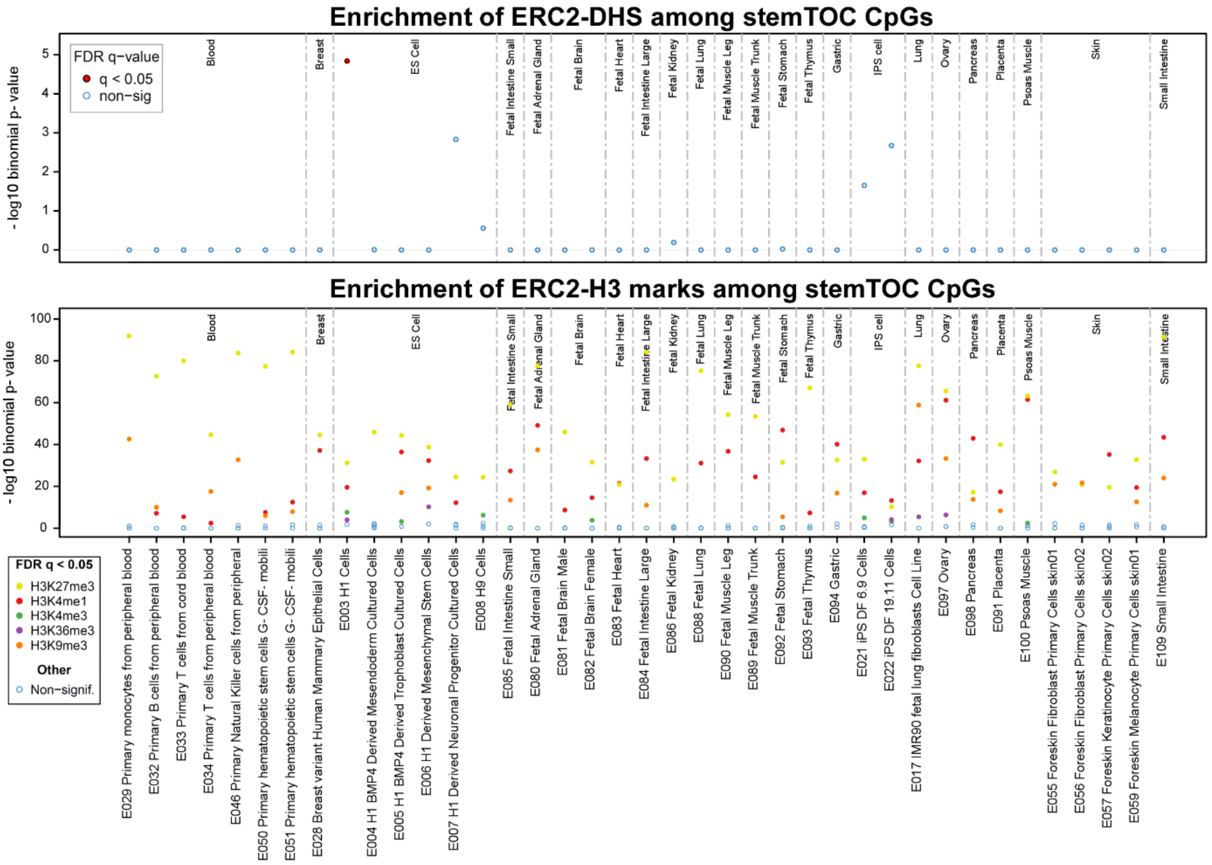


**Supplementary Figure 4a: eFORGE2 DHS and H3 enrichment analysis of stemTOC CpGs.** Top panel displays the -$\log_{10}$P-values of enrichment (y-axis) of DHSs from the eFORGE2 Binomial test among the 371 stemTOC CpGs. The enrichment is shown for DHSs as defined by different cell-types (x-axis). Lower panel is the corresponding enrichment plot for Histone H3 marks, as indicated. In both panels, colors indicates statistical significance at FDR<0.05.

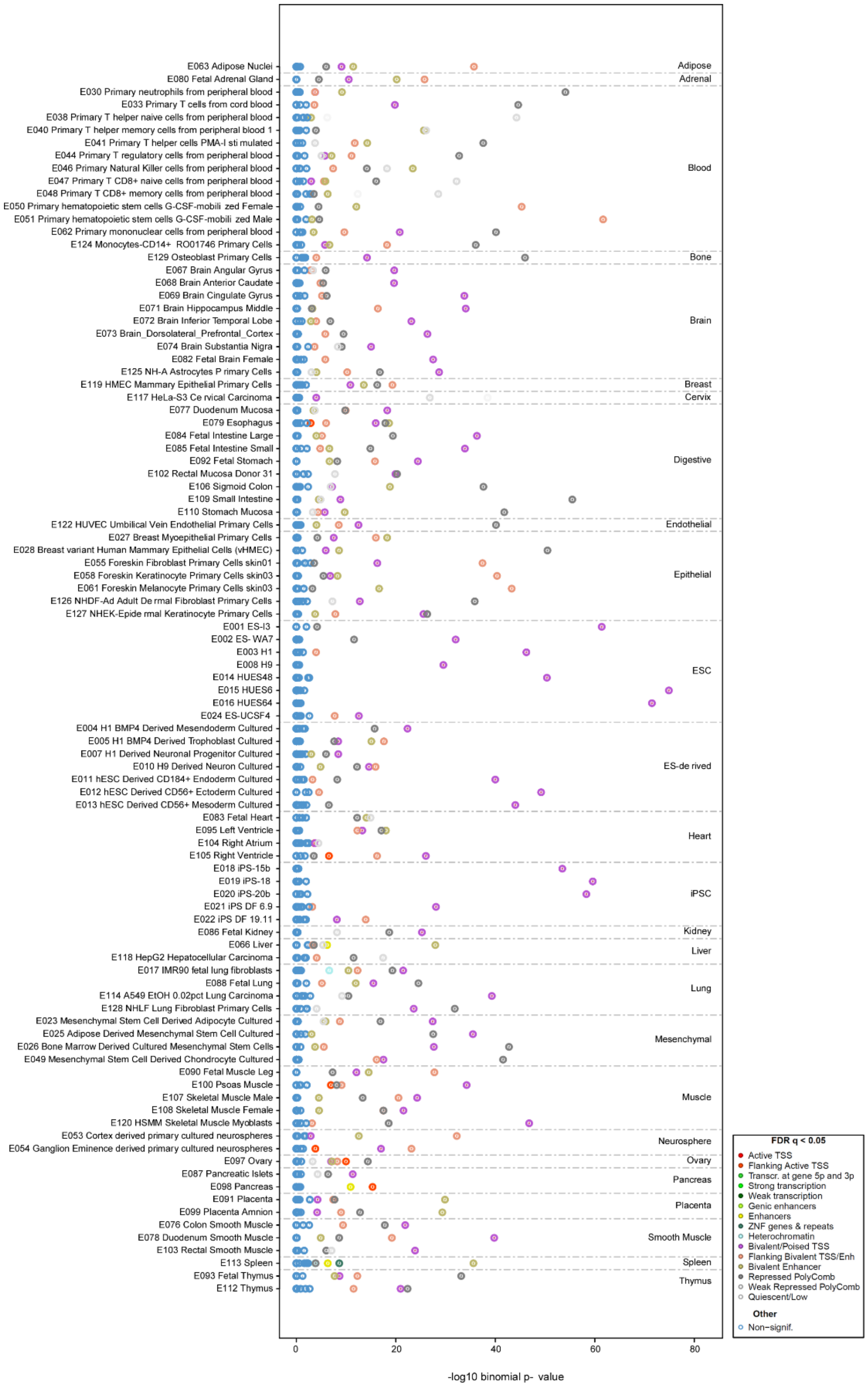


**Supplementary Figure 4b: eFORGE2 chromatin state enrichment analysis of stemTOC CpGs.** Panel displays the -$\log_{10}$P-values of enrichment (y-axis) of 15 chromatin states from the eFORGE2 Binomial test among the 371 stemTOC CpGs. The enrichment is shown for chromatin-states as defined by different cell-types (x-axis). Colors indicates statistical significance at FDR<0.05.

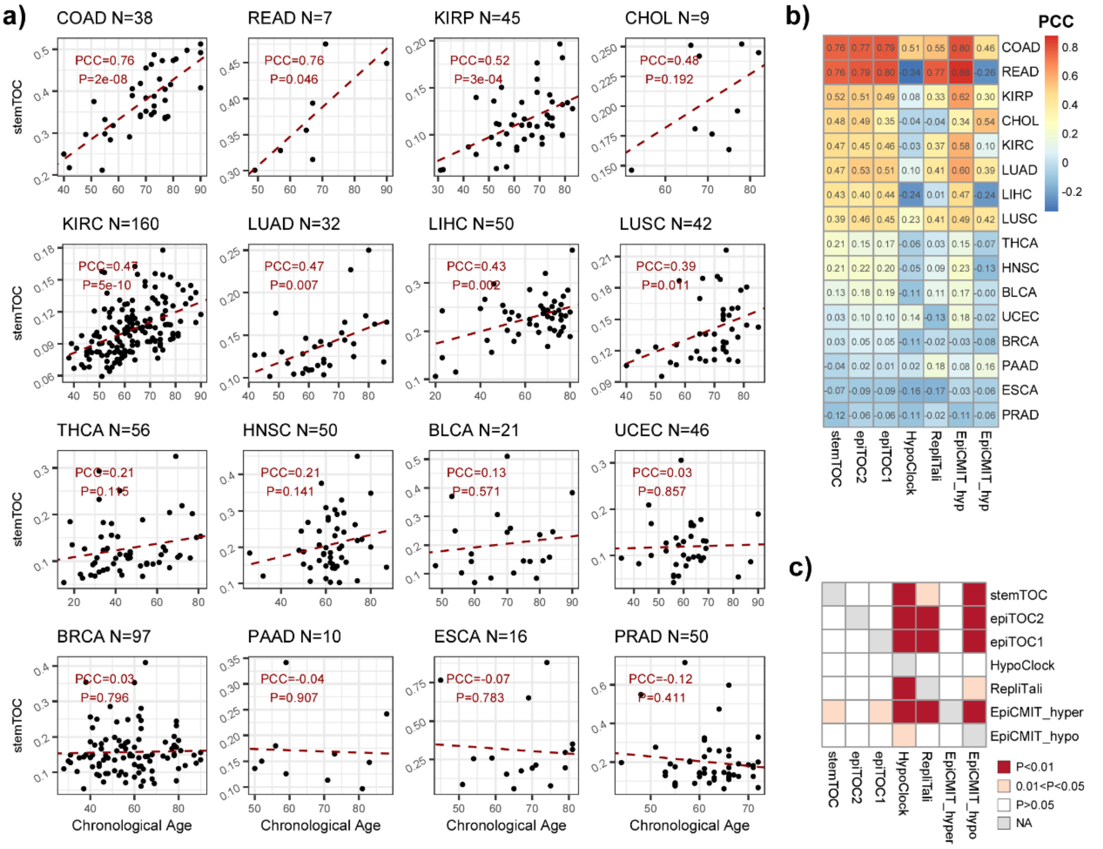


**Supplementary Figure 5: Correlation of mitotic-age with chronological age in normal-adjacent tissues from TCGA. a)** Scatterplots of the stemTOC mitotic age score (y-axis) against chronological age (x-axis) for 16 normal-adjacent tissue-types from TCGA. The number of samples in each age-group indicated below the plot. For each panel we provide the Pearson Correlation Coefficient (PCC) and P-value from a linear regression. **b)** Heatmap of corresponding PCC-values for each tissue-type and for 7 different mitotic clocks, as indicated. The PCC-values are provided in the heatmap. **c)** Heatmap displaying one-tailed paired Wilcoxon rank sum test P-values, comparing clocks to each other, in how well their mitotic age correlates with chronological age. Each row indicates how well the corresponding clock's mitotic age estimate correlates with chronological age as compared to the clock specified by the column. The paired Wilcoxon test is performed over the 16 tissue-types. RepliTali results are for the probes restricted to 450k beadarrays.

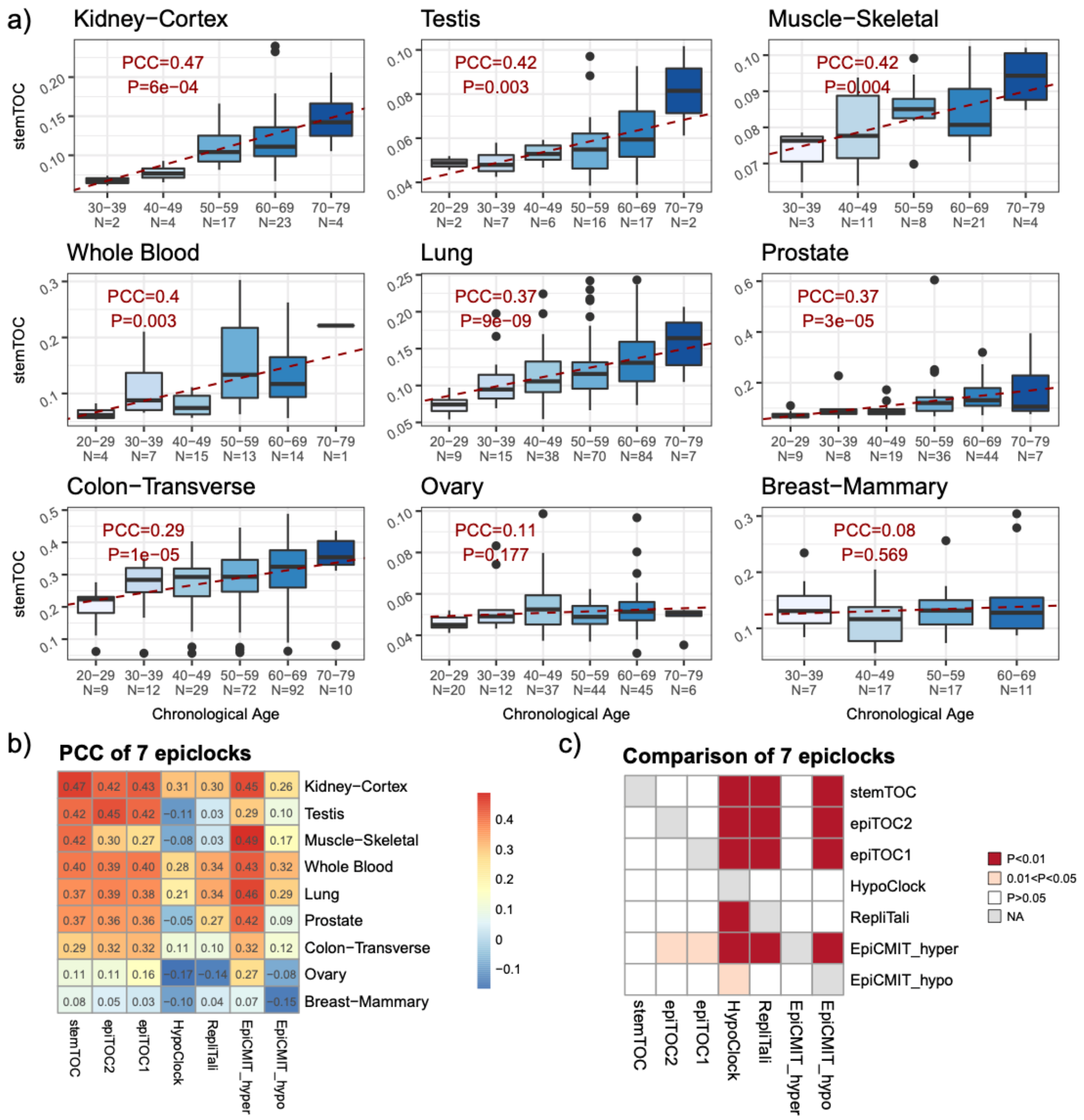


**Supplementary Figure 6: Correlation of mitotic-age with chronological age in normal tissues from eGTEX. a)** Boxplots of stemTOC mitotic age score (y-axis) against chronological age group (x-axis) for 9 normal tissue-types from eGTEX (EPIC data). The number of samples in each age-group indicated below the plot. For each panel we provide the Pearson Correlation Coefficient (PCC) and P-value from a linear regression. The line within each box denotes the median with the box itself defining the interquartile range and whiskers extending 1.5 times the interquartile range. **b)** Heatmap of corresponding PCC-values for each tissue-type and for 8 different mitotic clocks, as indicated. The PCC-values are provided in the heatmap. **c)** Heatmap displaying one-tailed paired Wilcoxon rank sum test P-values, comparing clocks to each other, in how well their mitotic age estimate correlates with chronological age. Each row indicates how well the corresponding clock's mitotic age estimate correlates with chronological age as compared to the clock specified by the column. The paired Wilcoxon test is performed over the 9 tissue-types.

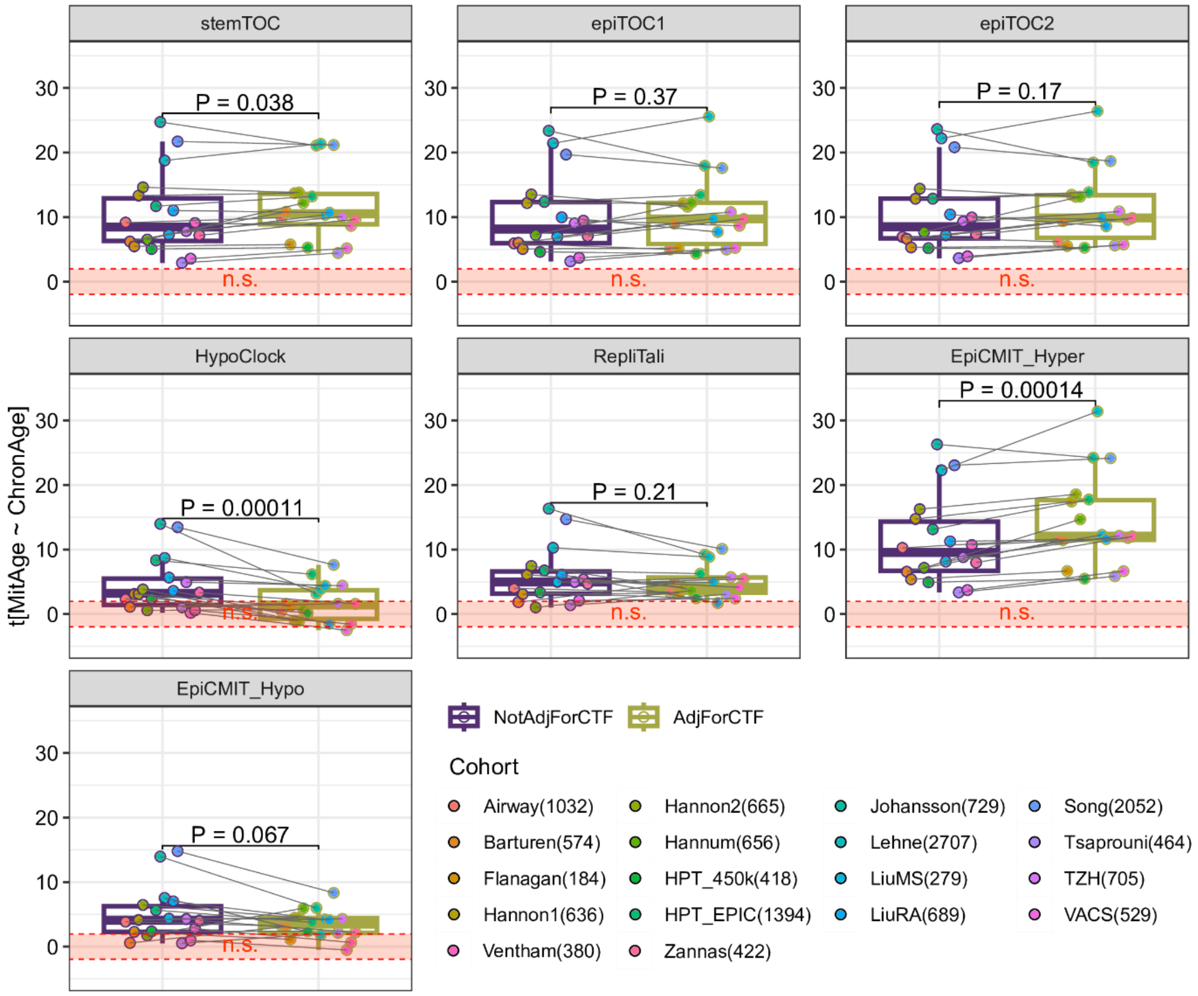


**Supplementary Figure 7: Effect of CTH on associations of mitotic age with chronological age in blood.** For each mitotic clock, boxplots compare the linear regression t-statistics of association between mitotic age and chronological age, not adjusting for immune cell-type fractions (NotAdjForCTF) and adjusting for 12 immune-cell type fractions (AdjForCTF). Each datapoint corresponds to one whole blood cohort, and there are a total of 18 whole blood cohorts, as indicated. The number of samples in each cohort is given in brackets after the cohort name. P-values derive from two-tailed paired Wilcoxon rank sum tests comparing the t-statistics before and after adjustment for CTFs. The line within each box denotes the median with the box itself defining the interquartile range and whiskers extending 1.5 times the interquartile range.

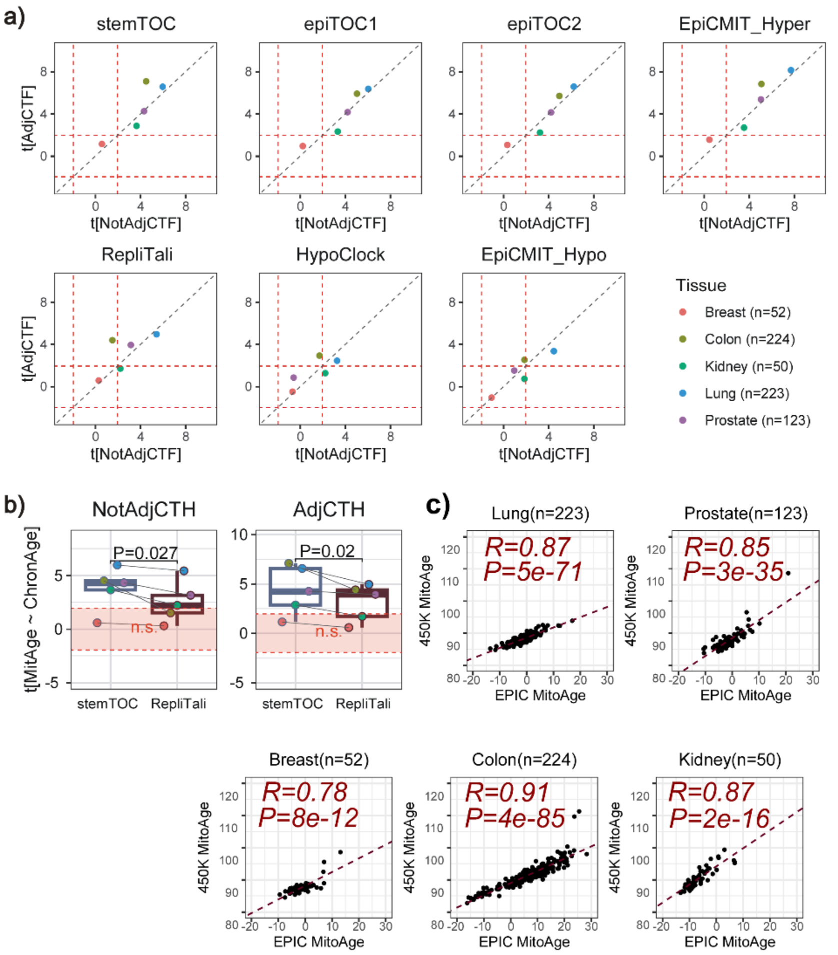


**Supplementary Figure 8: Effect of CTH on associations of mitotic age with chronological age in solid normal tissues from eGTEX (EPIC data). a)** Scatterplots of the linear regression t-statistics of association between mitotic age and chronological age, before (NotAdjCTF) and after (AdjCTF) adjustment for cell-type fractions. Each datapoint represent an eGTEX normal tissue EPIC dataset, for which the underlying cell-type fractions in the tissue could be estimated using our EpiSCORE DNAm-atlas algorithm. The red dashed-lines indicate the threshold of statistical significance (P=0.05). Number of samples in each dataset is given in brackets. **b)** Boxplots of the same linear regression t-statistics comparing stemTOC to RepliTali, not adjusting (left) and adjusting for CTH (right). P-values derive from a one-tailed paired t-test. The line within each box denotes the median with the box itself defining the interquartile range and whiskers extending 1.5 times the interquartile range. **c)** Scatterplots of RepliTali's Mitotic-Age in the 5 eGTEX normal tissue-datasets, computed using all available EPIC probes (x-axis) vs restricting to 450k probes only (y-axis). R-value and two-tailed correlation test P-value is given. Number of samples is given above each panel.

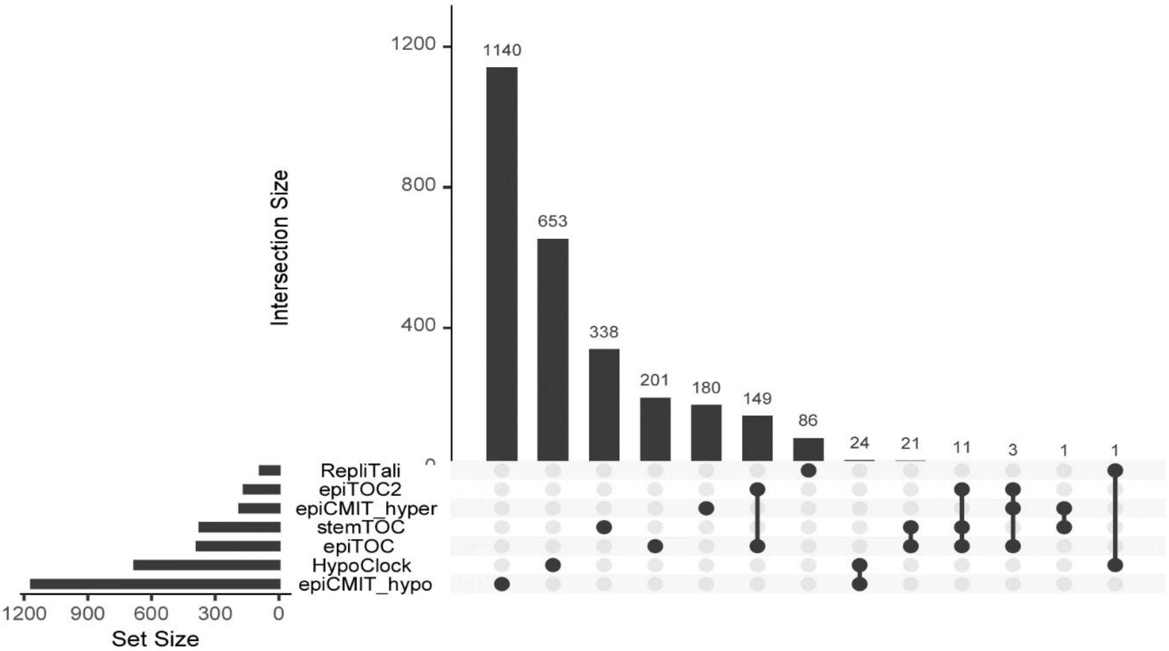


**Supplementary Figure 9: Upset plot displaying the CpG overlap of all 7 mitotic counters.** Barplots to on the bottom left indicate the number of CpGs making each mitotic-counter. Barplot on the top right not only indicate the number of CpGs in each counter/clock, but also the overlap between various counters. Of note, if a pair of clocks display zero overlap it is not shown. Thus, for instance, the overlap between RepliTali and all other clocks is zero, except for HypoClock with which it shares only 1 CpG site. stemTOC has zero overlap with all hypomethylated counters, has 1 CpG in common with epiCMIT-hyper, an overlap of 11 CpGs with epiTOC2, and an overlap of 21+11 CpGs with epiTOC.

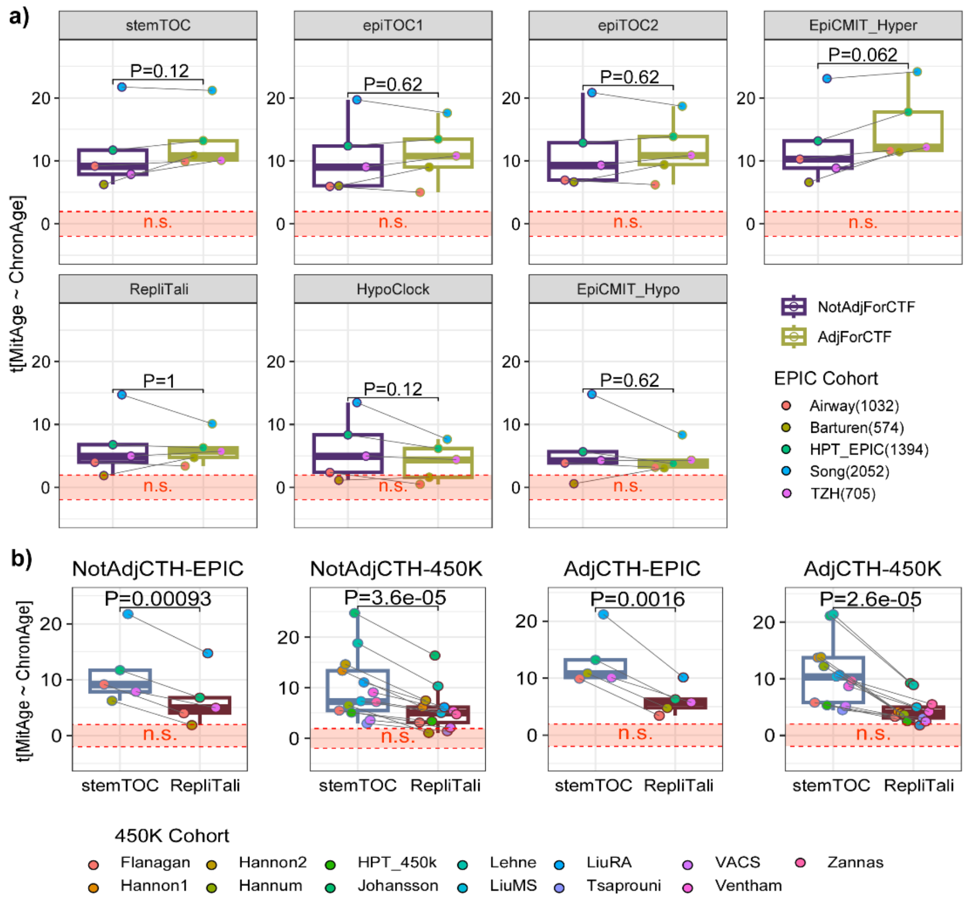


**Supplementary Figure 10: Effect of CTH on associations of mitotic age with chronological age in whole blood EPIC datasets. a)** For each mitotic clock, boxplots compare the linear regression t-statistics of association between mitotic age and chronological age, not adjusting for immune cell-type fractions (NotAdjForCTF) and adjusting for 12 immune-cell type fractions (AdjForCTF) in each of 5 whole blood EPIC DNAm datasets, as indicated. P-values derive from two-tailed paired Wilcoxon rank sum tests comparing the t-statistics before and after adjustment for CTFs. Number of samples in each EPIC cohort is given in brackets after the cohort name. **b)** A direct comparison of stemTOC to RepliTali, stratified by technology of DNAm dataset (EPIC or 450k) and by adjustment for cell-type heterogeneity (CTH) (NotAdjCTH or AdjCTH). Here the P-values are from a one-tailed paired t-test. The line within each box denotes the median with the box itself defining the interquartile range and whiskers extending 1.5 times the interquartile range. Number of samples of 450k cohorts: Flanagan (n=184), Hannon2 (n=665), HPT_450k (n=418), Lehne (n=2707), LiuRA (n=689), VACS (n=529), Zannas (n=422), Hannon1 (n=636), Hannum (n=656), Johansson (n=729), LiuMS (n=279), Tsaprouni (n=464), Ventham (n=380).

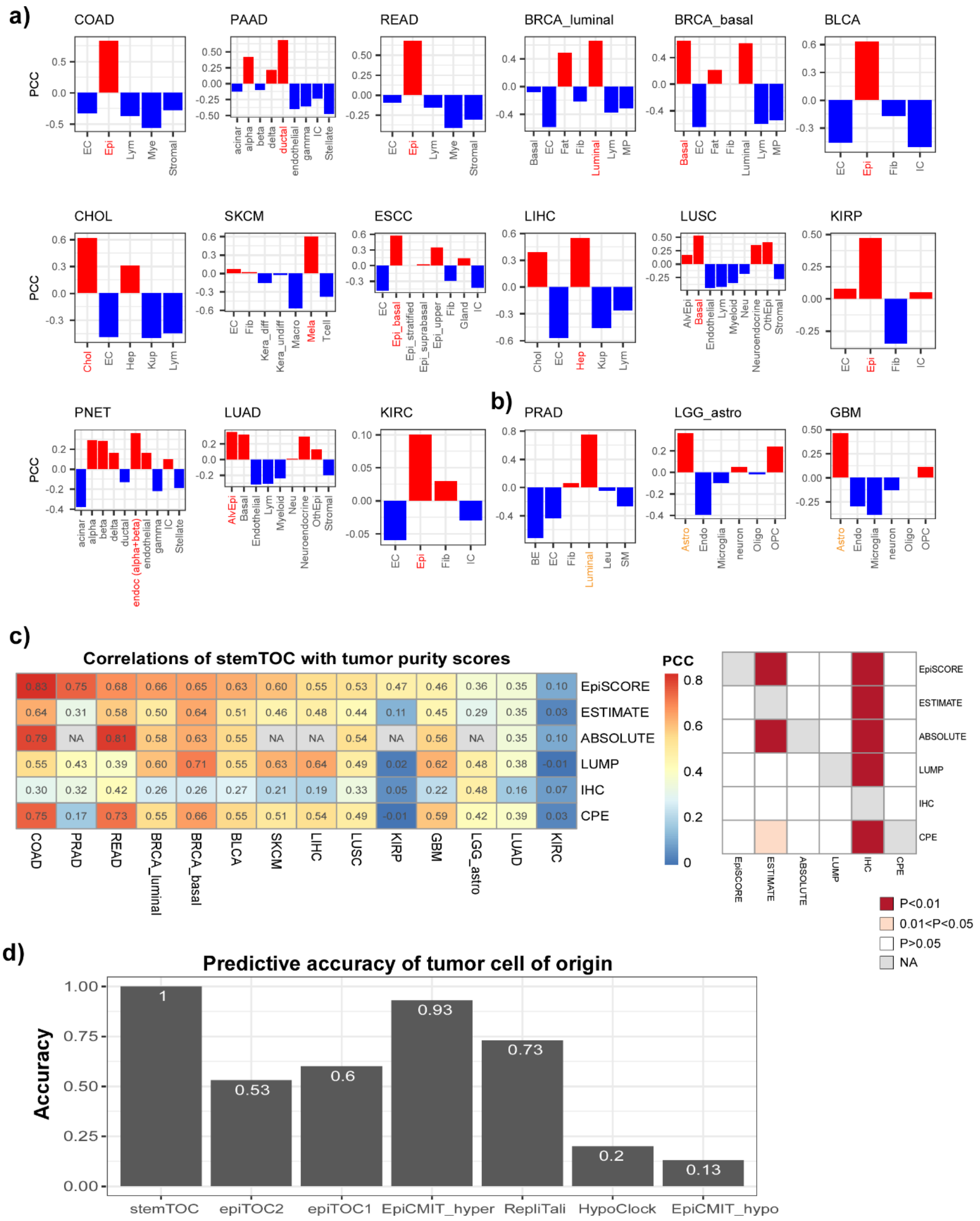


**Supplementary Figure 11: Predicting tumor cell-of-origin. a)** Barplots display Pearson Correlation Coefficient (PCC) between stemTOC's mitotic age and the corresponding cell-type fraction for each of 15 cancer-types where the tumor cell-of-origin (text in red) is fairly well established. **b)** As a) but for 3 TCGA cancer-types where the tumor cell-of- origin (text in orange) is less well established. **c)** Left: Heatmap of Pearson Correlation Coefficients (PCC) between stemTOC's mitotic age and tumor purity indices, including EpiSCORE's tumor cell of origin fraction, ESTIMATE, IHC, ABSOLUTE, CPE and LUMP. Right: Heatmap of one-tailed Wilcoxon rank sum test P-values comparing the PCCs obtained with each tumor purity method. A significant P-value for method "Y" on y-axis against method "X" on x-axis means significantly stronger PCCs with stemTOC's mitotic age for method "Y" compared to method "X". **d)** Overall prediction accuracy of 7 mitotic clocks for correctly predicting the tumor cell-of-origin, as assessed over the 15 cancer-types in a). RepliTali results are for the probes restricted to 450k beadarrays. Number of samples as in legend of Fig.3

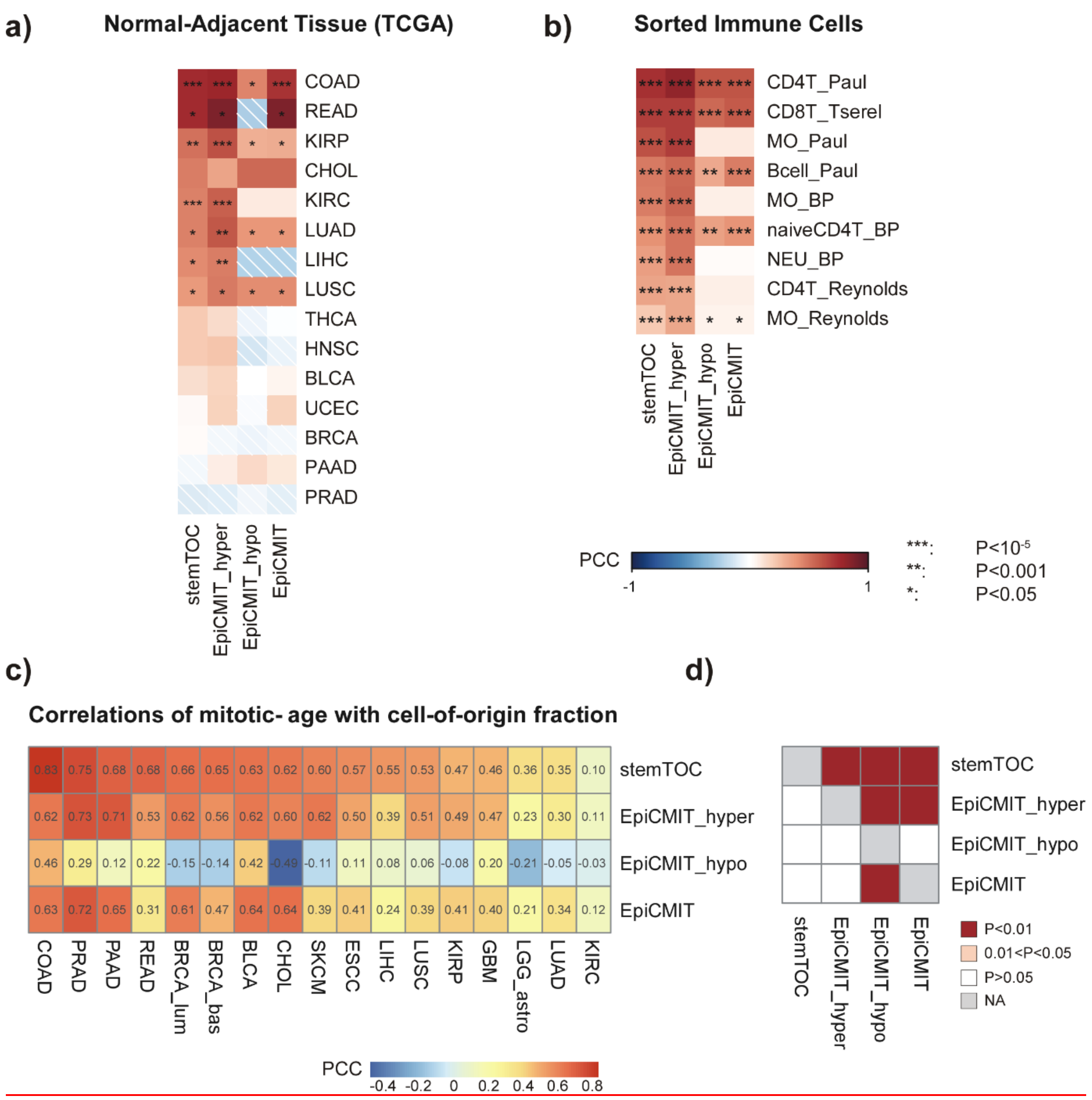


**Supplementary Figure 12: epiCMIT-hyper outperforms epiCMIT. a)** Pearson Correlation Coefficient (PCC) heatmap of correlations by 4 mitotic clocks (stemTOC, epiCMIT-hyper, epiCMIT-hypo, epiCMIT) with chronological in the normal-adjacent samples of TCGA cancer-types. **b)** As a) but for sorted immune-cell datasets. **c)** As a) but now for correlations of the mitotic clocks with tumor cell of origin fraction (as a proxy for tumor purity). **d)** Wilcoxon rank sum paired test one-tailed P-values comparing the PCC values displayed in c) of one clock to another. Convention is that a significant P-value means that the mitotic clock labelled by the row outperforms the one displayed in the column. Sample numbers in each cohort can be found in Fig.2 and Fig.2 legend.

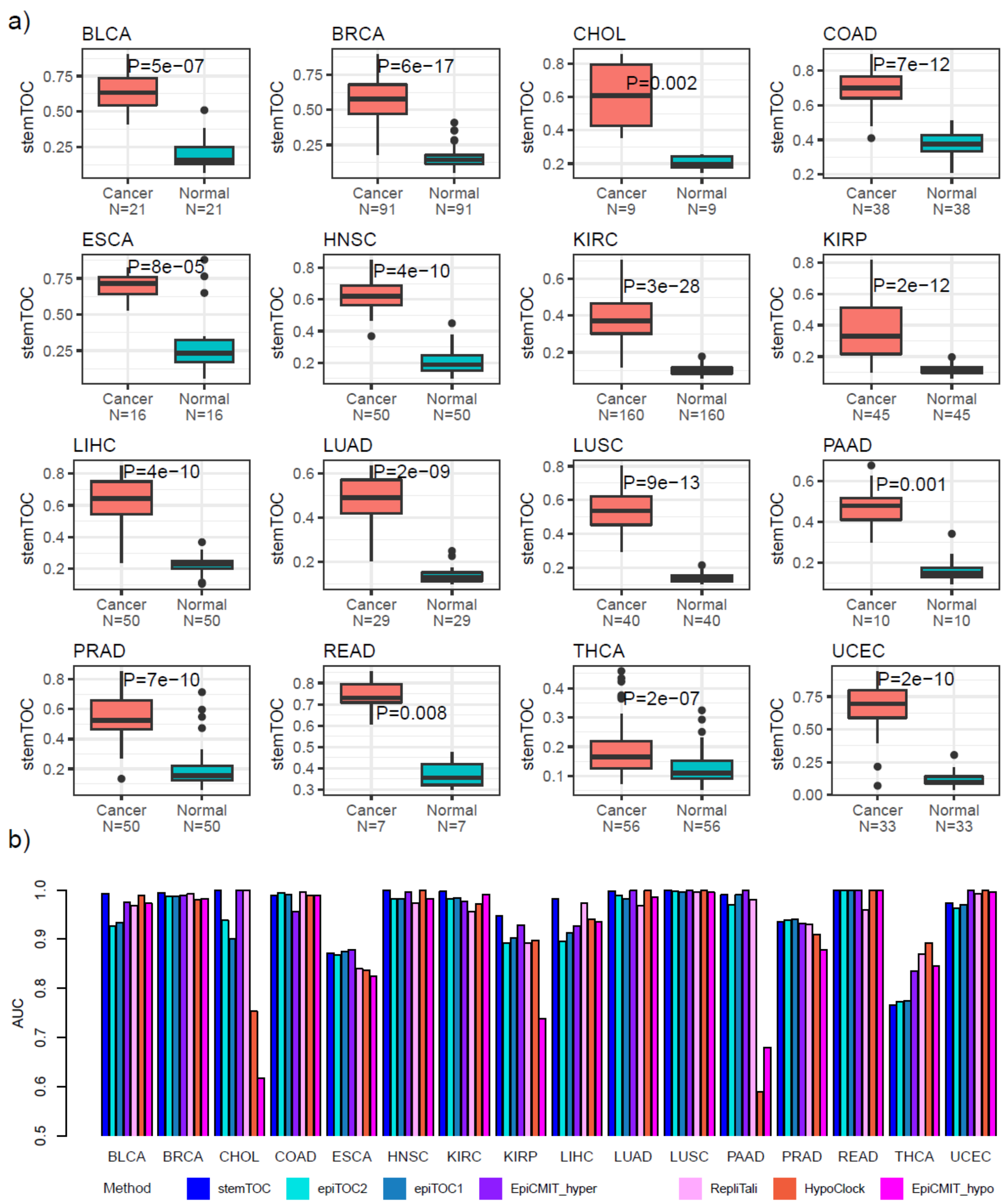


**Supplementary Figure 13: Mitotic counters discriminate cancer from normal-adjacent tissue. a)** Boxplots of stemTOC's mitotic age (y-axis) using only tumor normal-adjacent pairs for TCGA cancer types with sufficient numbers of normal samples. P-value derives from a one-tailed paired Wilcoxon rank sum test. Number of samples in each groups is given below boxplot. The line within each box denotes the median with the box itself defining the interquartile range and whiskers extending 1.5 times the interquartile range. **b)** Corresponding AUC-values for all mitotic clocks. RepliTali results are for the probes restricted to 450k beadarrays.

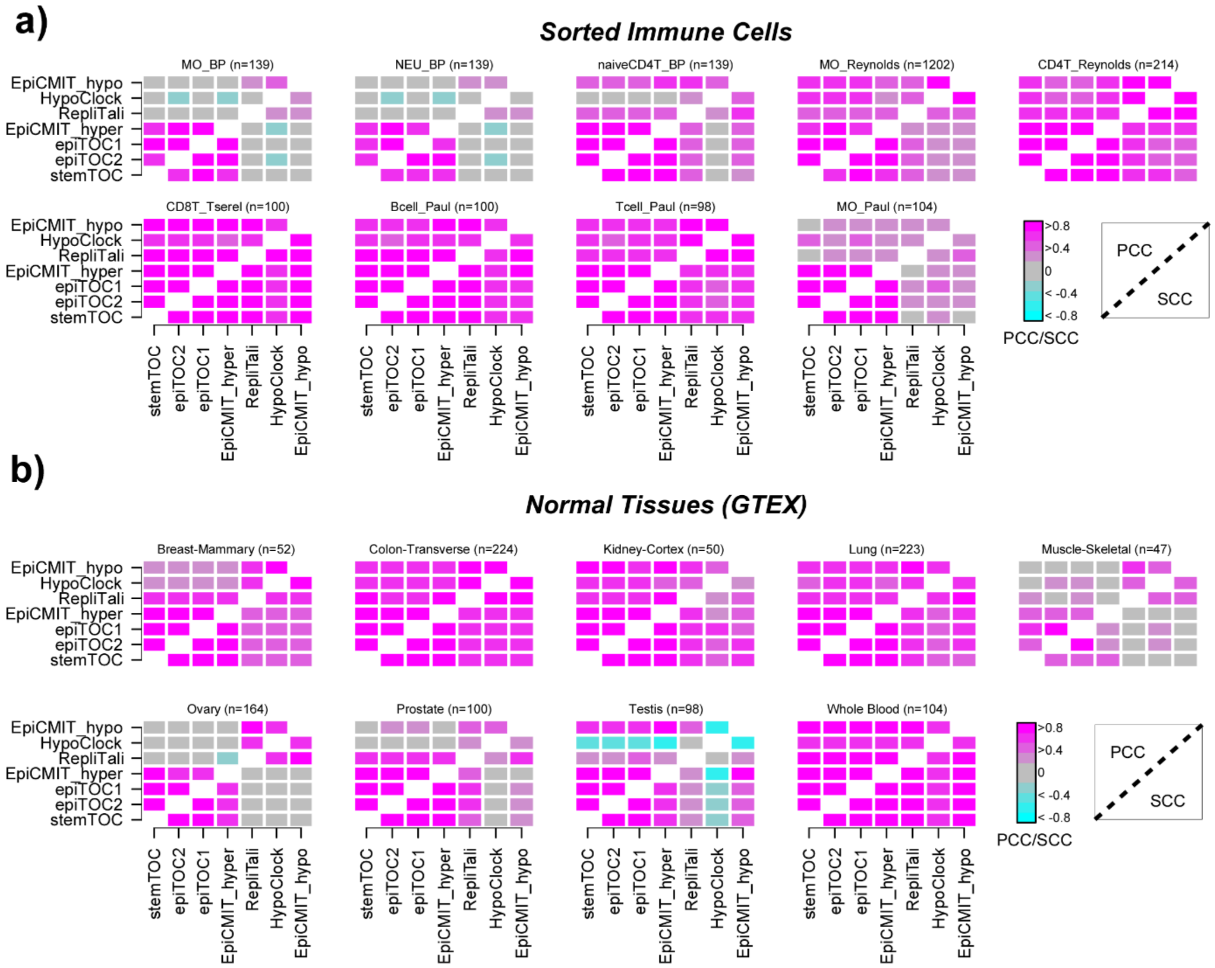


**Supplementary Figure 14: Correlation heatmaps of mitotic counters in sorted immune cells and normal tissues of GTEX. a)** For each dataset profiling sorted immune cells we display a heatmap of Pearson (upper triangular part, PCC) and Spearman (lower triangular part, SCC) correlation coefficients between each pair of mitotic counters. The cell-type and number of samples in each dataset are given above each heatmap. MO=monocyte, NEU=neutrophil, Tcell=CD4+ T-cell. CD4T=CD4+ T-cell, CD8T=CD8+ T-cell, naiveCD4T=naïve CD4+ T-cell. RepliTali results are for the probes restricted to 450k beadarrays. **b)** As a) but for the normal-tissue datasets from GTEX (EPIC data).

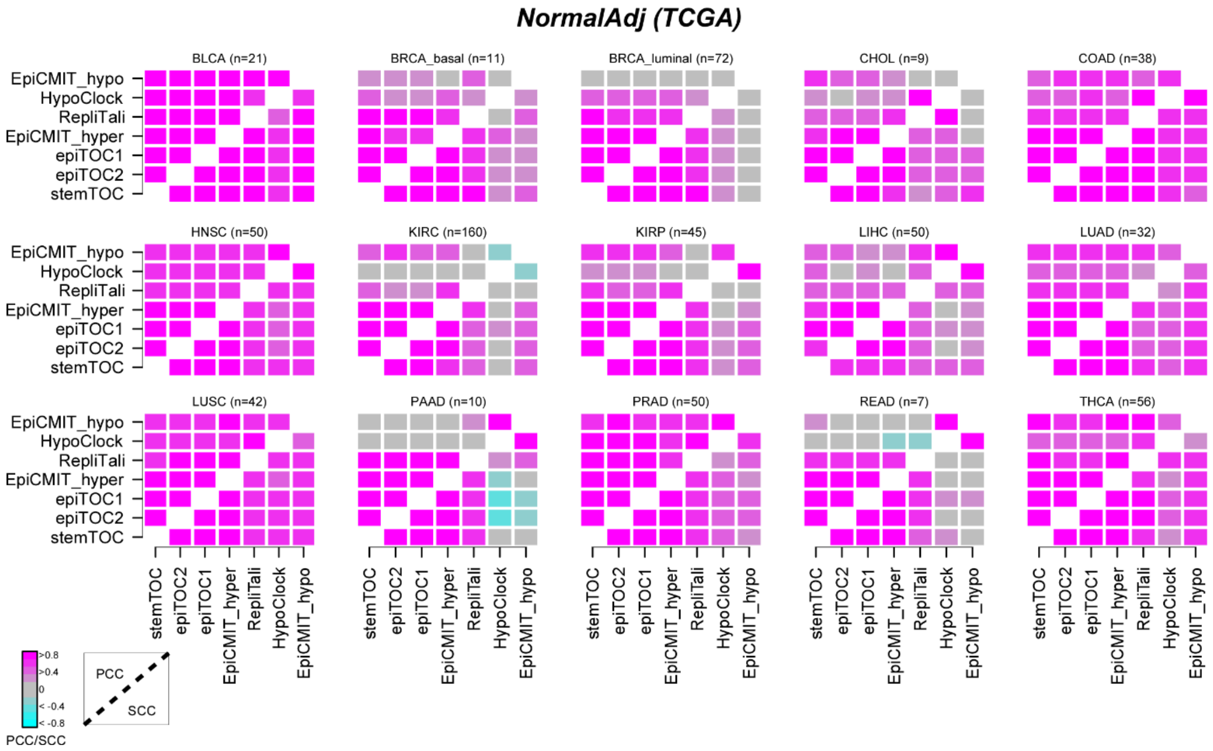


**Supplementary Figure 15: Correlation heatmaps of mitotic counters in the normal-adjacent samples from the TCGA.** For each set of normal-adjacent samples we display a heatmap of Pearson (upper triangular part, PCC) and Spearman (lower triangular part, SCC) correlation coefficients between each pair of mitotic counters. The cancer-type matched to the normal-adjacent tissue and number of normal-adjacent samples in each dataset are given above each heatmap. RepliTali results are for the probes restricted to 450k beadarrays.

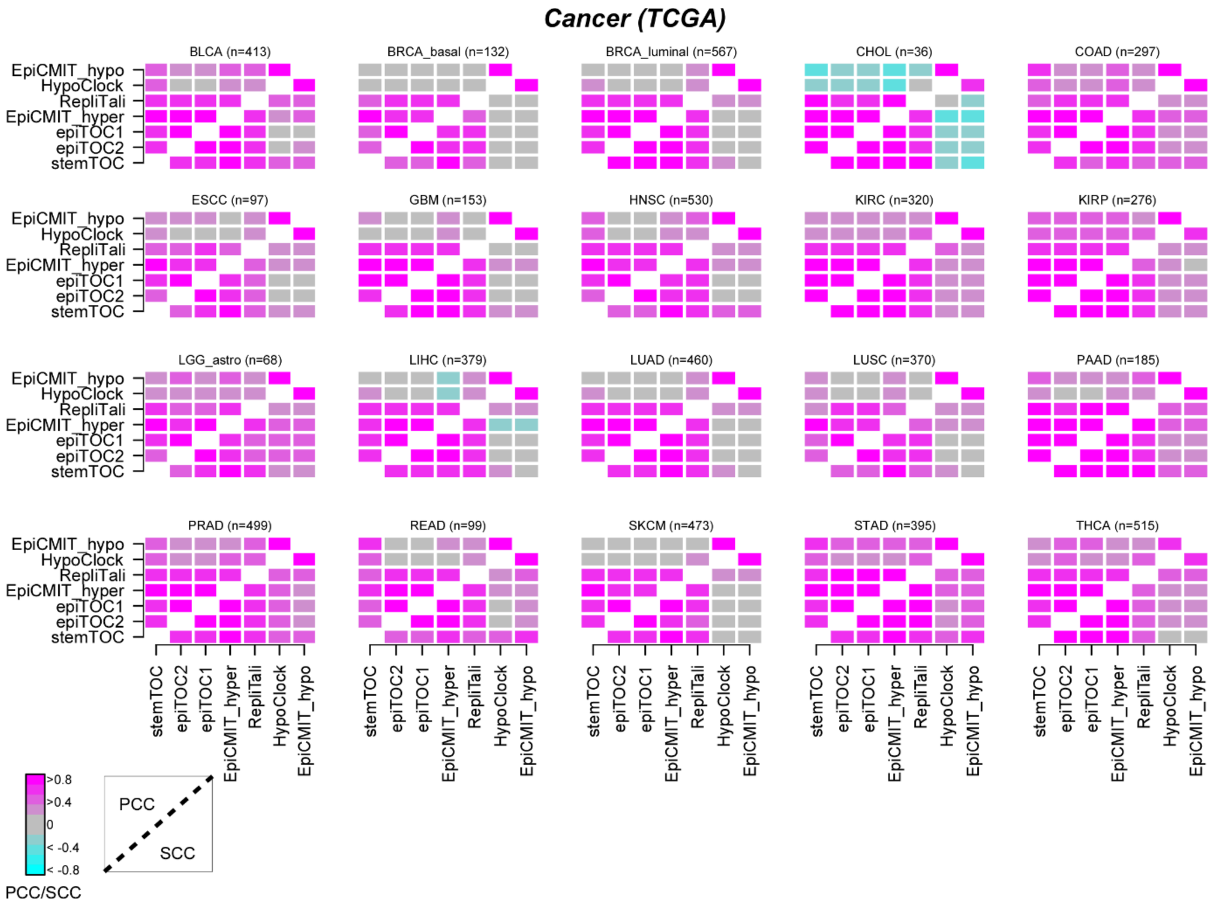


**Supplementary Figure 16: Correlation heatmaps of mitotic counters in the tumor samples from the TCGA.** For each TCGA cancer-type we display a heatmap of Pearson (upper triangular part, PCC) and Spearman (lower triangular part, SCC) correlation coefficients between each pair of mitotic counters. The cancer-type and number of tumor samples in each dataset are given above each heatmap. RepliTali results are for the probes restricted to 450k beadarrays.

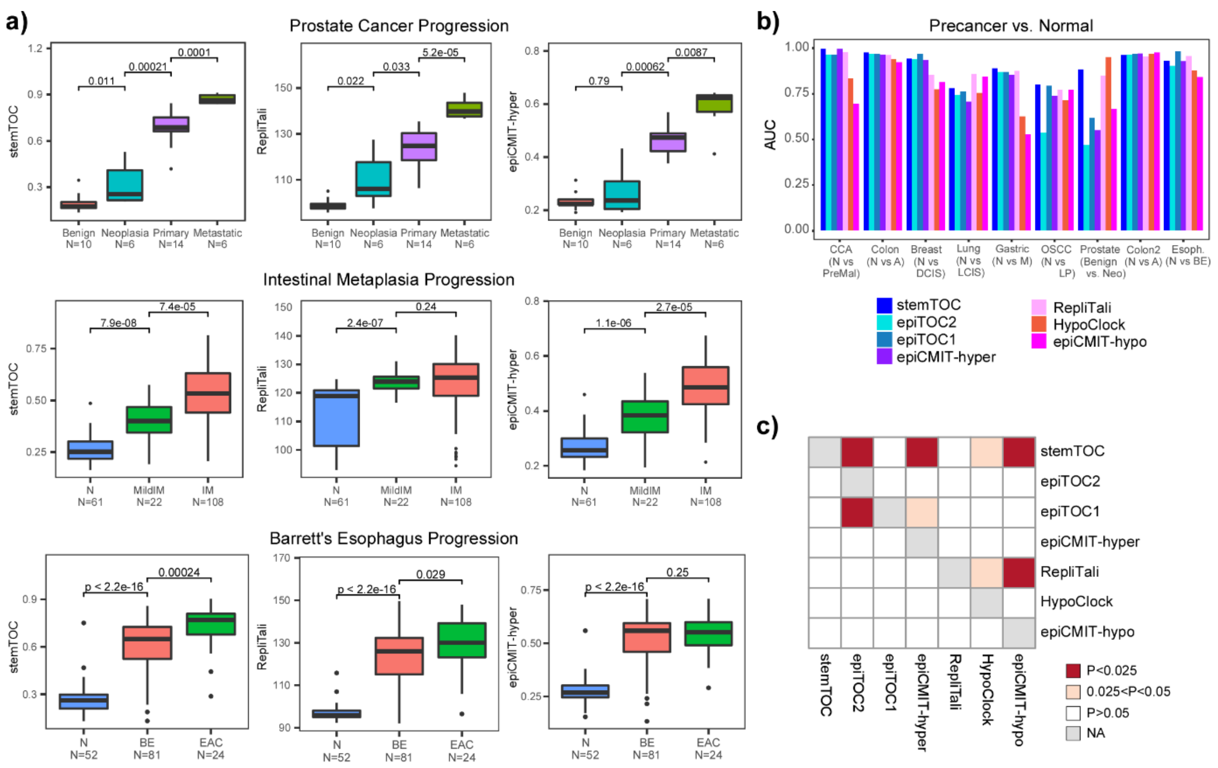


**Supplementary Figure 17: Benchmarking of stemTOC in precancerous conditions . a)** Mitotic-age estimates for stemTOC, RepliTali and epiCMIT-hyper in three different DNAm datasets encompassing normal healthy tissue and precancerous lesions, including prostate cancer progression (Benign, Neoplasia, Primary and Metastasis), progression of intestinal metaplasia (N=normal, MildIM=mild intestinal metaplasia, IM=intestinal metaplasia) and esophageal adenocarcinoma (N=normal healthy squamous, BE=Barrett's Esophagus, EAC=esophageal adenocarcinoma). Number of samples in each group is indicated. Number of samples in each group is shown below boxplot. We provide one-tailed P-values from a Wilcoxon rank sum test comparing successive stages. The line within each box denotes the median with the box itself defining the interquartile range and whiskers extending 1.5 times the interquartile range. **b)** Barplot displaying the AUC from the Wilcoxon test comparing normal-healthy to normal at-risk groups across a total 9 different DNAm datasets, and for 7 different mitotic clocks. Number of samples is as in Fig.4a. **c)** Heatmap displaying one-tailed paired Wilcoxon rank sum test P-values, comparing clocks to each other, in how well their mitotic age distinguished normal-healthy from normal at-cancer-risk tissue. Each row indicates how well the corresponding clock's mitotic age estimate performs in relation to the clock specified by the column. The paired Wilcoxon test is performed over the 9 datasets. RepliTali results in 450k datasets are for the probes restricted to 450k beadarrays.

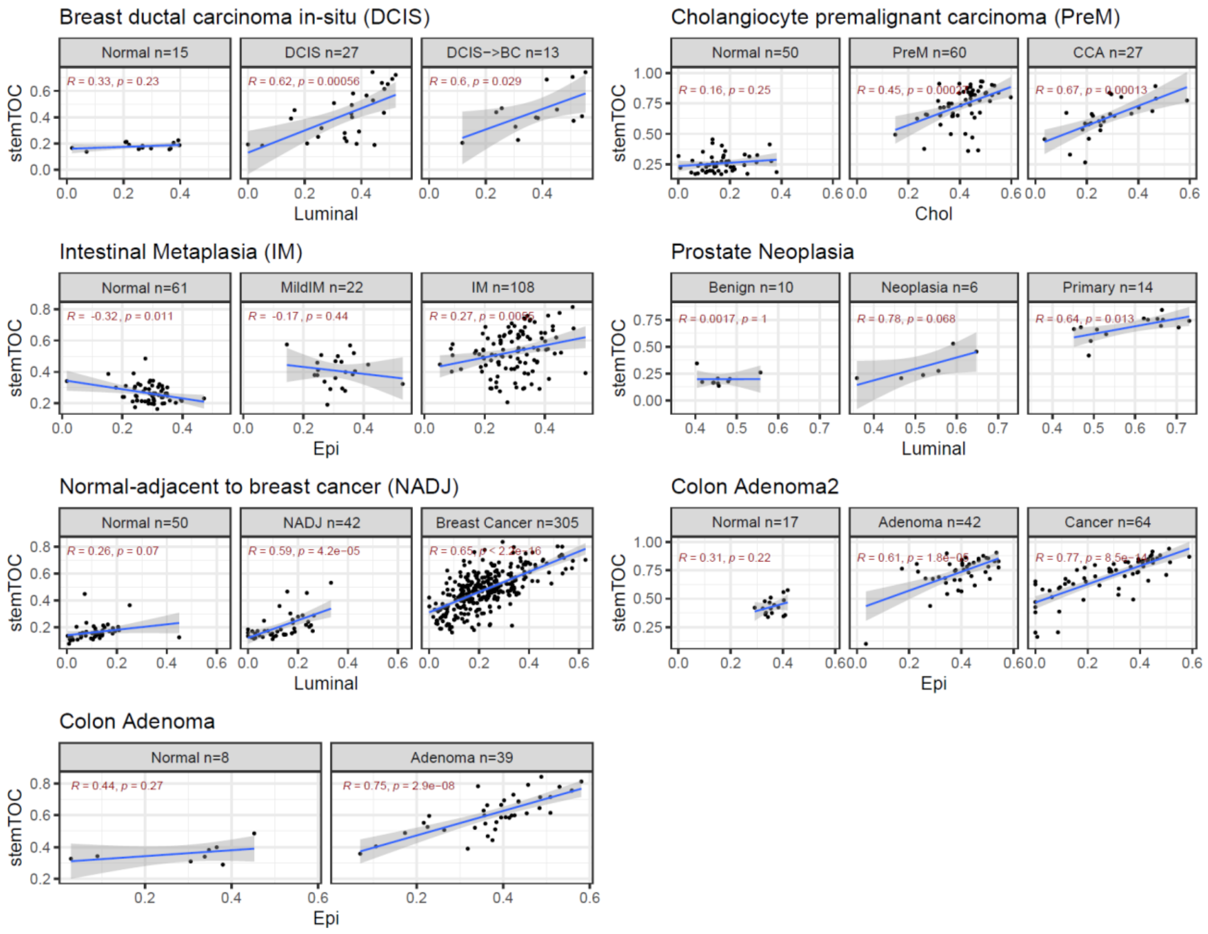


**Supplementary Figure 18: Correlations of stemTOC's mitotic age with tumor cell-of-origin fraction in precancerous lesions.** Scatterplots of stemTOC's mitotic age (y-axis) vs the EpiSCORE-estimated fraction of the presumed cell-of-origin for a number of Illumina EPIC/450k DNAm datasets profiling histologically normal and precancerous lesions, as indicated. In each scatterplot we give the R-value and P-value from a linear regression. The number of samples is indicated above the plot. Regression line with standard error interval is shown.

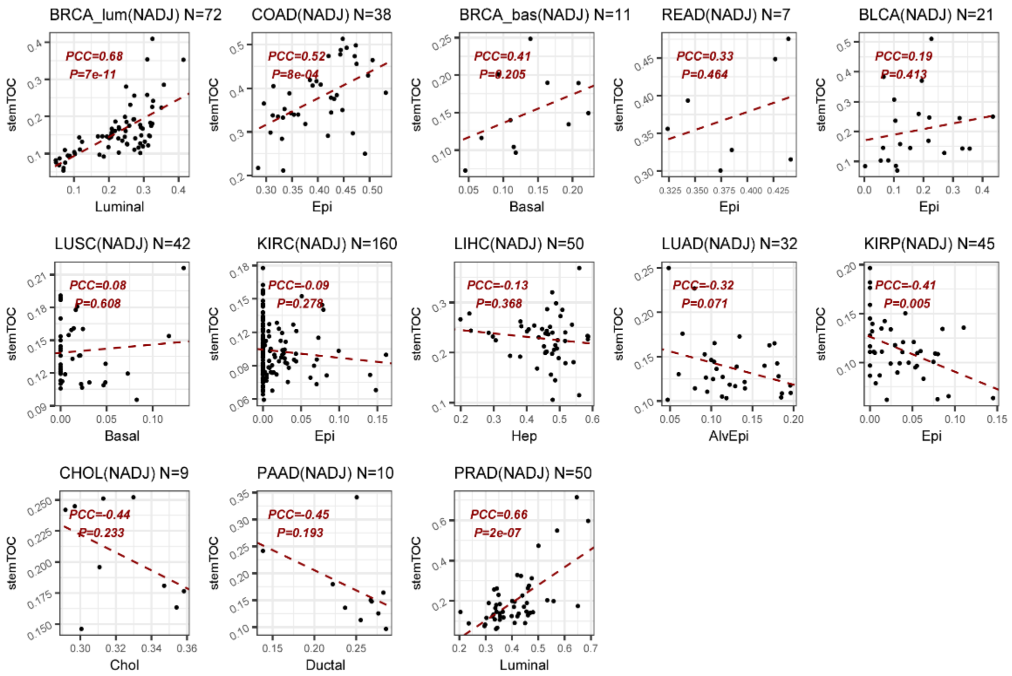


**Supplementary Figure 19: Correlations of stemTOC's mitotic age with tumor cell-of-origin fraction in the normal-adjacent tissues of the TCGA.** Scatterplots of stemTOC's mitotic age (y-axis) vs the EpiSCORE-estimated fraction of the presumed cell-of-origin in normal-adjacent (NADJ) tissue datasets from the TCGA. Only those with reasonable numbers of NADJ samples were included. In each scatterplot we give the R-value and P-value from a linear regression. The number of samples is indicated above the plot. Regression line is shown.

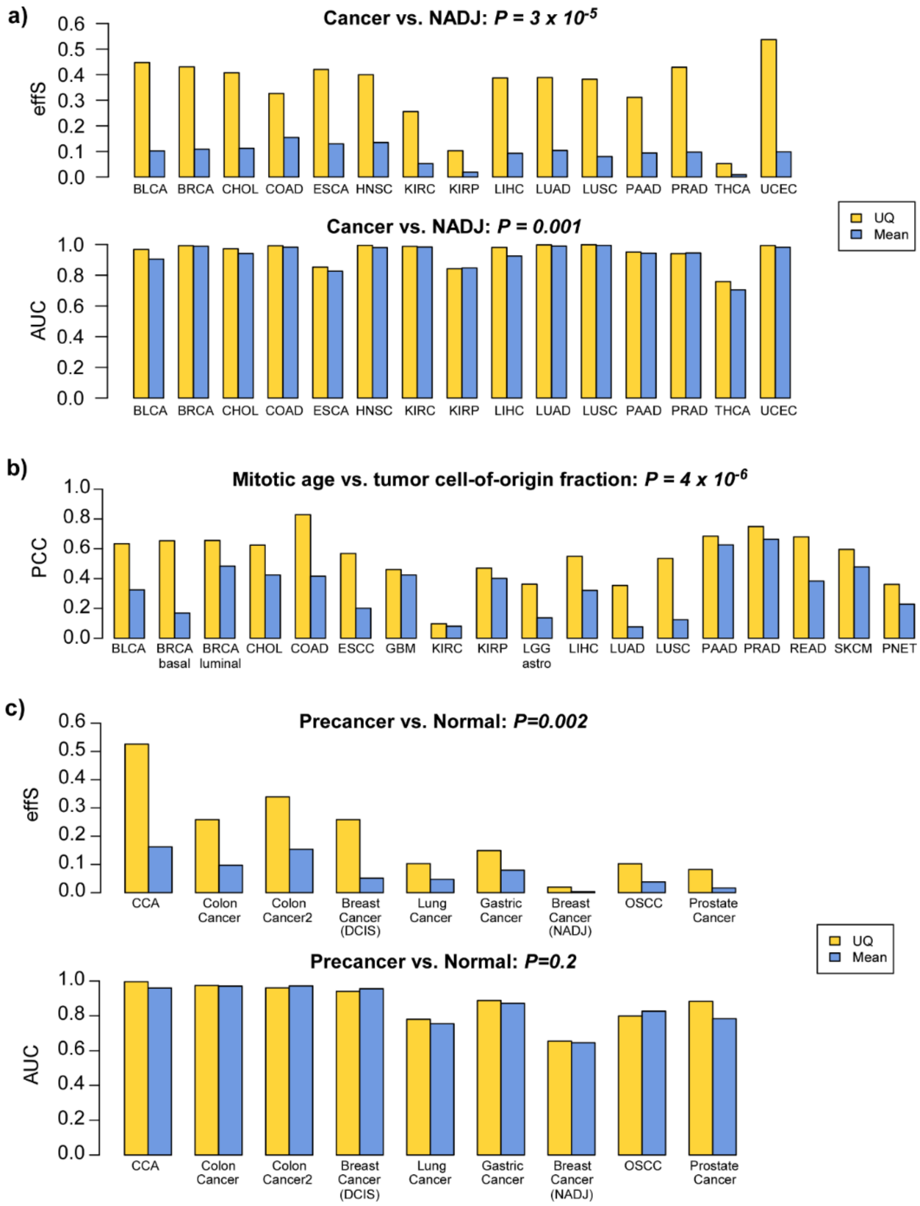


**Supplementary Figure 20: Accounting for stochasticity improves associations of mitotic age. a)** Barplots depict the effect size (effS) of the average mitotic age difference between cancer and normal-adjacent tissue for TCGA cancer-types, and with the mitotic age computed using a 95% upper quantile (UQ, stemTOC) or the mean (Mean) over the 371 mitotic-CpGs. Lower barplots display the corresponding AUCs. For both barplots, we provide the one-tailed P-value from a paired Wilcoxon test comparing UQ to mean. **b)** Barplots depicts the Pearson Correlation Coefficient (PCC) between mitotic age and tumor cell of origin fraction across TCGA cancer-types. We provide the one-tailed P-value from a paired Wilcoxon test comparing the PCCs from using UQ (stemTOC) to those using the mean. **c)** Upper barplots depict the effect size (effS) of the average mitotic age difference between normal-healthy and normal “at-cancer-risk” tissue for various cancer-types. Barplots below display the corresponding AUCs. For both sets of barplots, we provide the one-tailed P-value from a paired Wilcoxon test comparing UQ to mean.

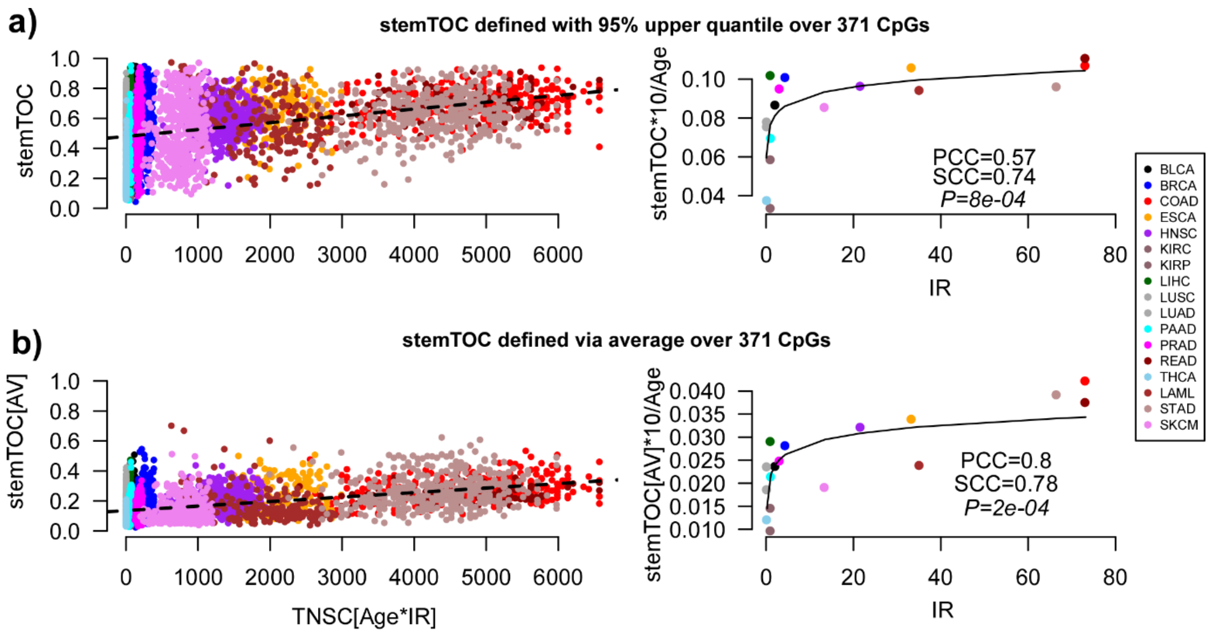


**Supplementary Figure 21: Saturation effect of stemTOC is not dependent on taking an upper quantile. a) Left:** Scatterplot of stemTOC (defined using the 95% upper quantile) (y-axis) against the estimated total number of stem-cell divisions (Age*intrinsic annual rate of stem-cell division of the corresponding normal-tissue (IR), x-axis) for >7000 cancer samples from 17 TCGA cancer-types. Only cancer-types with an IR estimate in normal tissue were used. **Right:** Median value of age-adjusted stemTOC value (multiplied by 10 to reflect change over a decade) for each cancer-type vs the annual intrinsic rate of stem-cell division of the corresponding normal tissue-type. Both Pearson (PCC) and Spearman (SCC) correlation coefficients are given. P-value tests for significance of SCC. Fitted line is that of a best fit among linear and log-linear models. **b)** As a), but defining stemTOC as the average DNAm over the 371 stemTOC CpGs. Sample sizes of each cancer cohort given as in Fig.3.

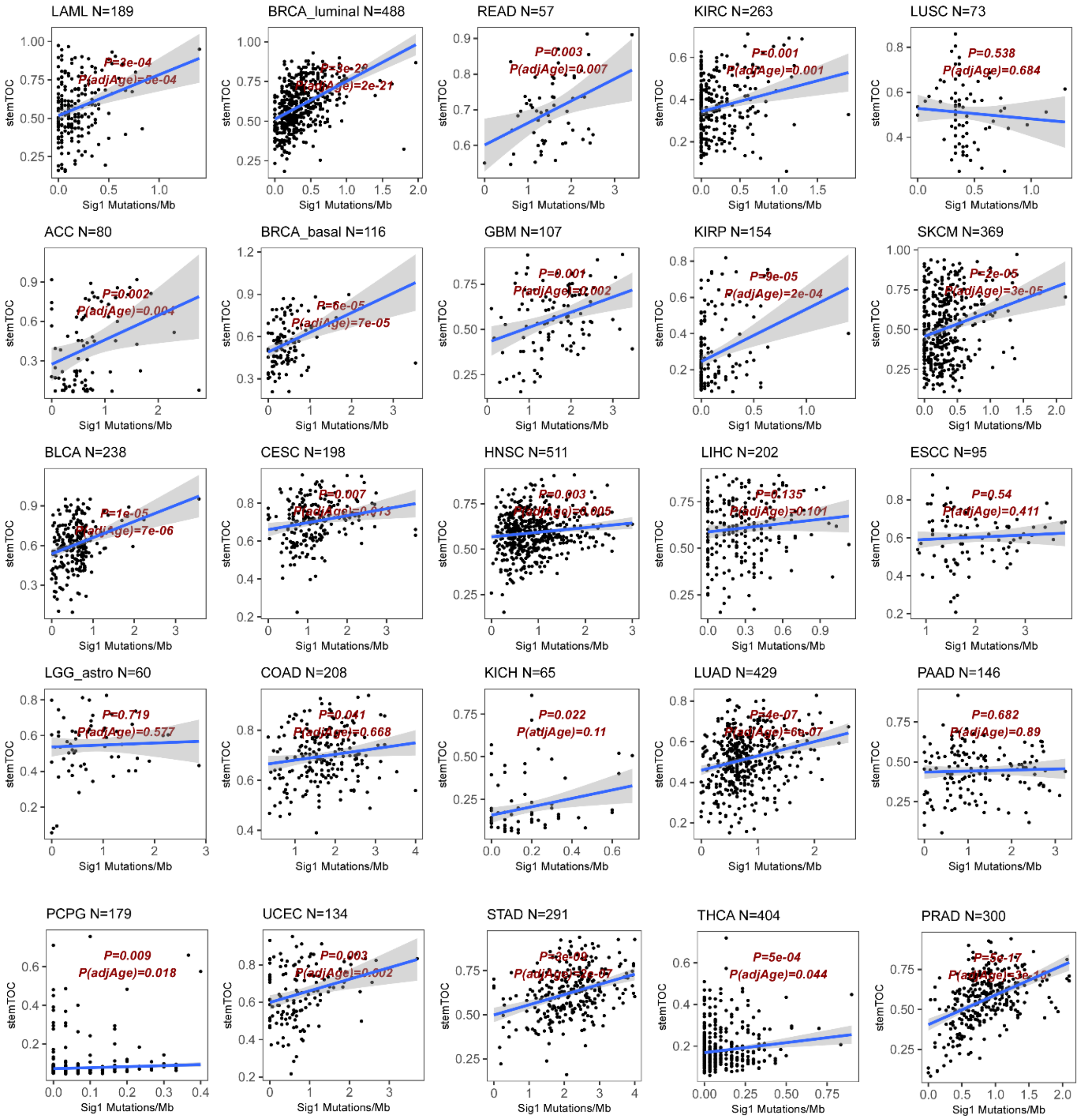


**Supplementary Figure 22: stemTOC vs MS1 in individual TCGA cancer-types.** Scatterplot of stemTOC's mitotic age vs MS1 mutational load (mutations per Mb) for 25 TCGA cancer-types, as shown. Number of cancer samples is shown above plots. In each panel, we display the P-values of a linear regression of stemTOC's mitotic age vs the mutational load, adjusting for age (P(adjAge)) and not adjusting for age (P). Regression line with standard error interval is shown.

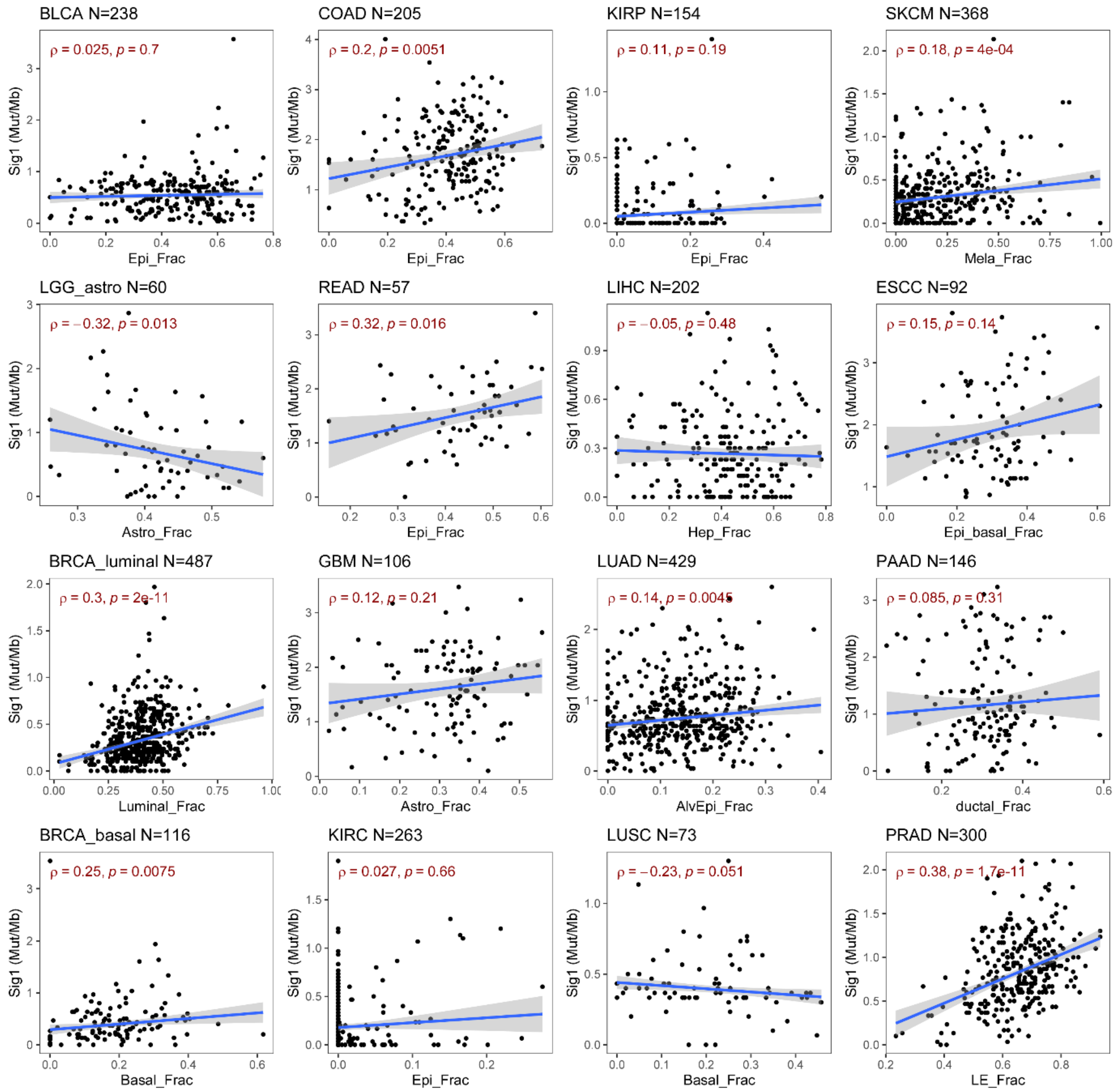


**Supplementary Figure 23: MS1 load vs tumor cell of origin fraction.** Scatterplot of MS1 mutational load (mutations per Mb) vs tumor cell of origin fraction (as estimated using the EpiSCORE algorithm), for 16 TCGA cancer-types, as shown. Number of cancer samples is shown above plots. In each panel, we display Spearman's correlation coefficient and P-value. Regression line with standard error interval is shown.

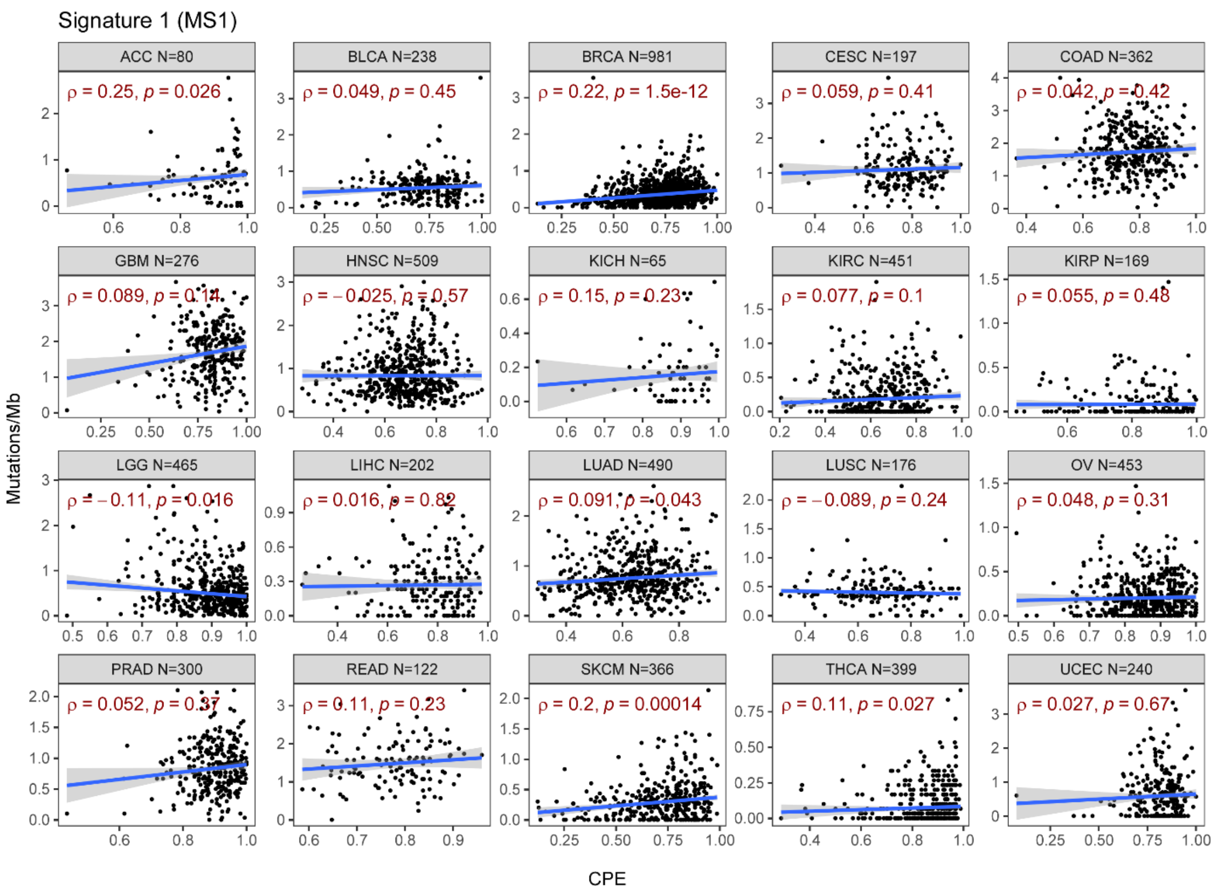


**Supplementary Figure S24: MS1 load does not generally correlate with tumor purity.** Scatterplot of MS1 mutational load (mutations per Mb) vs tumor purity (as estimated using the CPE algorithm), for 20 TCGA cancer-types, as shown. Number of cancer samples is shown above plots. In each panel, we display the Spearman Correlation Coefficient and P-value. Regression line with standard error interval is shown.
